# CD95L concatemers highlight difference in the manner CD95-mediated apoptotic and non-apoptotic pathways are triggered

**DOI:** 10.1101/2023.06.22.546070

**Authors:** Eden Lebrault, Christelle Oblet, Keerthi Kurma, Nicolas Levoin, Mickael Jean, Pierre Vacher, Patrick Legembre

## Abstract

To better understand the stoichiometry of CD95L required to trigger apoptotic and non-apoptotic signals, we generated several CD95L concatemers from dimer to hexamer conjugated *via* a flexible link (GGGGS)_2_. These ligands reveal that although the hexameric structure is the best stoichiometry to trigger cell death, a dimer is sufficient to induce the apoptotic response in CD95-sensitive Jurkat cells. Interestingly, only trimeric and hexameric forms can implement a potent Ca^2+^ response, suggesting that while CD95 aggregation controls the implementation of the apoptotic signal, both aggregation and conformation are required to implement the Ca^2+^ pathway.

## Introduction

CD95L (FasL/CD178), a member of the tumor necrosis factor (TNF) family, contributes to immune homeostasis and anti-tumor and anti-infectious responses (1). CD95L is a type II transmembrane cytokine mainly expressed by activated T cells and NK cells to kill infected and transformed CD95-expressing cells. CD95L contains an extracellular stalk region cleaved by matrix metalloproteinases (MMP) and a disintegrin and metalloproteases (ADAMs) and a C-terminal TNF homology domain (THD) (2). After cleavage by metalloproteases, the soluble extracellular region of CD95L (sCD95L) exhibits a homo-trimeric stoichiometry (3,4). On the other hand, the transmembrane CD95L (mCD95L) can be capped at the plasma membrane of activated T cells and NK cells to bind and aggregate its receptor, CD95 (also known as Fas). Binding of mCD95L to CD95 promotes the recruitment of the adaptor protein FADD (Fas Associated Death Domain) through homotypic interactions between their respective death domains (DDs) (5). FADD in turn aggregates the initiator caspase-8 to form a CD95/FADD/caspase-8 complex, called death-inducing signaling complex (DISC), that can initiate the apoptotic program (6,7). sCD95L fails to trigger cell death but its binding to CD95 favors the recruitment of PLCγ1 at the plasma membrane (4,8), where the lipase cleaves phosphatidylinositol-4,5-bisphosphate into diacylglycerol and inositol triphosphate (IP3). IP3 in turn activates IP3 receptors (IP3Rs) on endoplasmic reticulum (ER) to release Ca^2+^. This reduction of Ca^2+^ in ER lumen is sensed by STIM1 and STIM2, which traffic to the plasma membrane to activate Orai channels, allowing Ca^2+^ influx from the extracellular space (9-12), a molecular mechanism designated store-operated calcium entry (SOCE).

A potential explanation for the failure of sCD95L to induce cell death came from its stoichiometry, since the metalloprotease-cleaved ligand exhibits an homo-trimeric structure (4,13,14) and the minimal stoichiometry required to induce apoptosis has been suggested to be an hexamer (13). This explanation is in agreement with the fact that a signaling threshold exists to activate CD95, and while low aggregation level induces non-apoptotic signaling pathways, higher aggregation triggers cell death (15-17). To decipher whether the only parameter controlling the implementation of the apoptotic *versus* non-apoptotic signaling (*i*.*e*., calcium response) pathways was the CD95L stoichiometry, we designed a set of CD95L concatemers and assessed their biological effects.

## Results and discussion

### Generation of CD95L concatemers

Modeling and structural studies point out that soluble CD95L is a compact, non-covalently-linked homotrimer (3, 4) whose binding to its receptor seems to occur by the insertion of the ligand into three individual CD95 receptors to initiate cell signaling pathways. Soluble CD95L exhibits a “jelly-roll” fold of β sheets (18), and its crystal structure was not complete since the stalk region encompassing amino acid residues 103 to 143 was missing. Because this stalk region has been suggested to modulate the aggregation level and/or conformation of the soluble CD95L (sCD95L)(19,20), we reconstituted this region by performing molecular dynamics simulations with the CD95L homo-trimer (Fig.1A and 1B). The final state of the reconstituted extracellular CD95L trimeric model highlighted that the N- and C-terminal regions of sCD95L were at proximity and thereby, we wondered whether this feature could be used to generate concatemers by covalently linking CD95L monomers with a flexible linker (*i*.*e*., GGGGS; Fig.1C).

**Figure 1.**
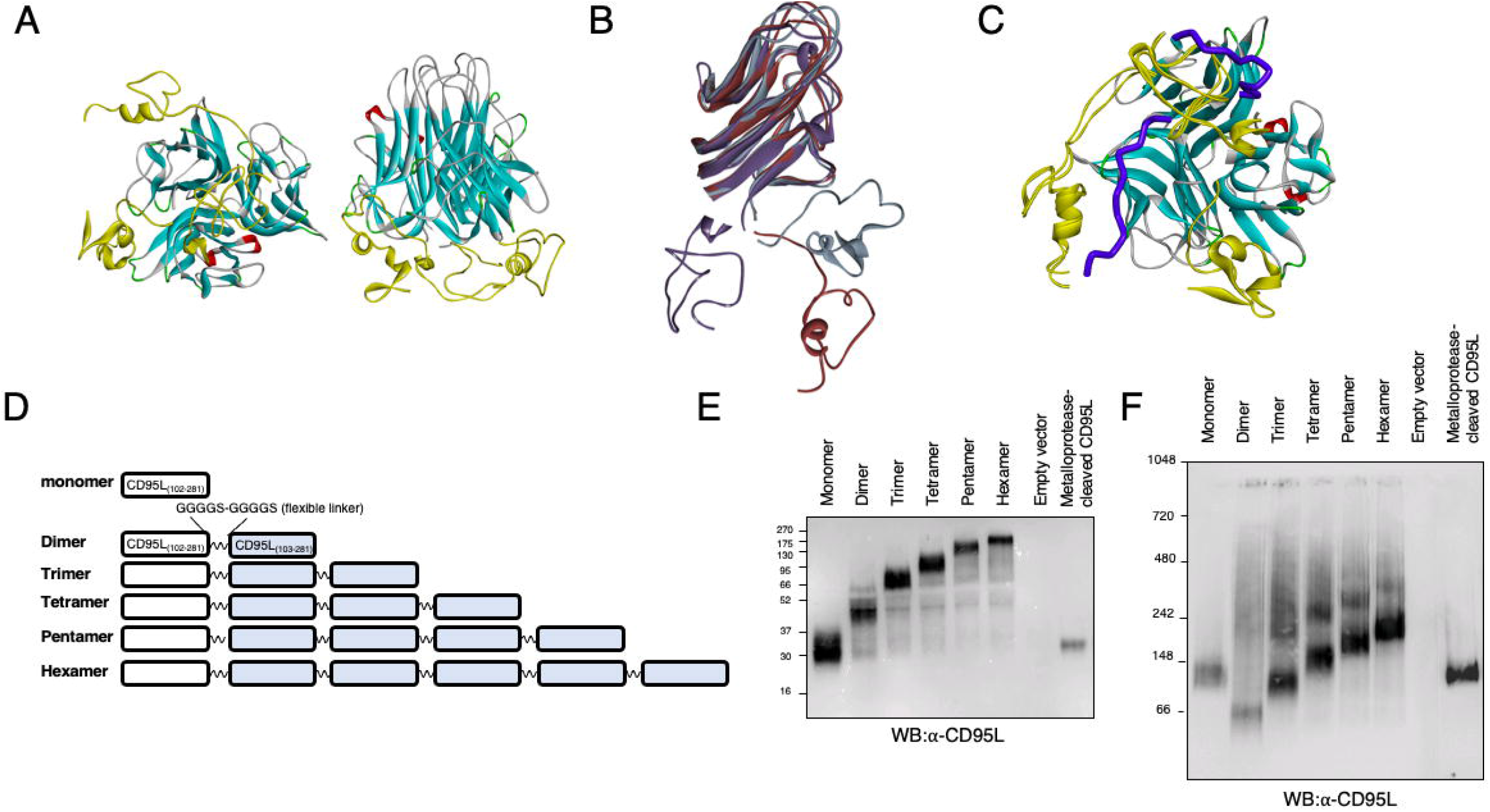
Generation of CD95L concatemers. **A**. The amino acid residues 103 to 143 were missing in the crystal structure. Their structure was built through extensive molecular dynamic simulations. The final state of the reconstituted CD95L trimeric model with Nt region in yellow is depicted. **B**. Addition of the amino acid residues 103 to 143 to the crystalized CD95L did not affect the experimental 3D structure of the protomer. **C**. Based on the reconstituted CD95L trimeric model showed in Figure A, a molecular modeling approach was applied to predict the minimal repetition of the flexible linker GGGGS (in blue) to connect covalently three CD95L protomers without affecting the global structure of the crystalized homo-trimeric CD95L. **D**. Schematic representation of the indicated concatemers. **E**. The CD95L concatemer-encoding pcDNA3 vectors and empty or human full length CD95L-encoding pLVX-IRES-tdTomato vectors were transfected in HEK/293T cells and supernatants were harvested after 7 days. After ultracentrifugation to eliminate exosomes, supernatants were resolved in reducing and denaturing conditions (SDS-PAGE). Indicated immunoblotting was performed. **F**. The molecular weights of the different CD95L were assessed in native condition using BN-PAGE method.

Because molecular modeling of the CD95L homotrimer predicted that the length and flexibility of a GGGGS dimer (*i*.*e*., (GGGGS)_2_) would be sufficient to connect CD95L monomers without affecting their global conformation (Fig.1C), we generated concatemers using this linker (Fig.1D). All constructs were transfected in the mammalian cell line HEK/293T and produced for in a serum-low medium (*i*.*e*., OPTI-MEM). Next, exosomes were eliminated by ultracentrifugation and the secreted ligands were analyzed in reducing and denaturating conditions by SDS-PAGE (Fig.1E) and in native condition using BN-PAGE (Fig.1F). Denaturing and reducing conditions (*i*.*e*., SDS-PAGE) confirmed that both the secreted CD95L monomer and the metalloprotease-cleaved CD95L possessed similar molecular weights (MW) around 30 kDa (Fig.1E). Although this MW was superior to the theoretical molecular weight 20.4 kDa according to Expasy website, this difference was in agreement with the N-glycosylation of CD95L in mammalian cells (21). In native conditions, the recombinant soluble CD95L (a.a. 102 to 281) and its metalloprotease-cleaved counterpart exhibited a molecular weight below 148 kDa suggesting a homotrimeric or tetrameric stoichiometry (Fig.1F). Of note, in native conditions, the dimeric concatemer migrated at 66 kDa, below its monomer counterpart suggesting that dimers failed to self-associate and form either tetramers or hexamers (Fig.1F). We also noticed that for trimeric, tetrameric, pentameric and hexameric concatemers, a faint band appeared at a higher molecular weight suggesting the formation of self-aggregate (Fig.1F).

### CD95L concatemers and cell death

The ability of each construct to trigger cell death was assessed by exposing the CD95-sensitive Jurkat T-cell to different concentrations of the concatemers (Fig.2A). Except the monomer, all concatemers exerted a cytotoxic effect (Fig.2A). Interestingly, while an hexamer of CD95L has been reported to be the minimal stoichiometry necessary to trigger cell death (13), we observed that the dimeric concatemer killed Jurkat cells in a similar manner to trimer, tetramer and pentamer counterparts (EC_50_ of 0.17 μg/mL for the dimer, *vs* 0.15 for the trimer, 0.08 for the tetramer and 0.13 for the pentamer). Nonetheless, the most efficient stoichiometry to kill remained the hexamer (13), since this ligand killed 4 to 9 fold more efficiently Jurkat cells as compared to the other concatemers (EC_50_ of 0.02 μg/mL). These findings suggested that although a dimeric CD95L was sufficient to trigger cell death, the CD95-mediated apoptotic signal possess a threshold level that can be reached by the hexameric complex.

**Figure 2.**
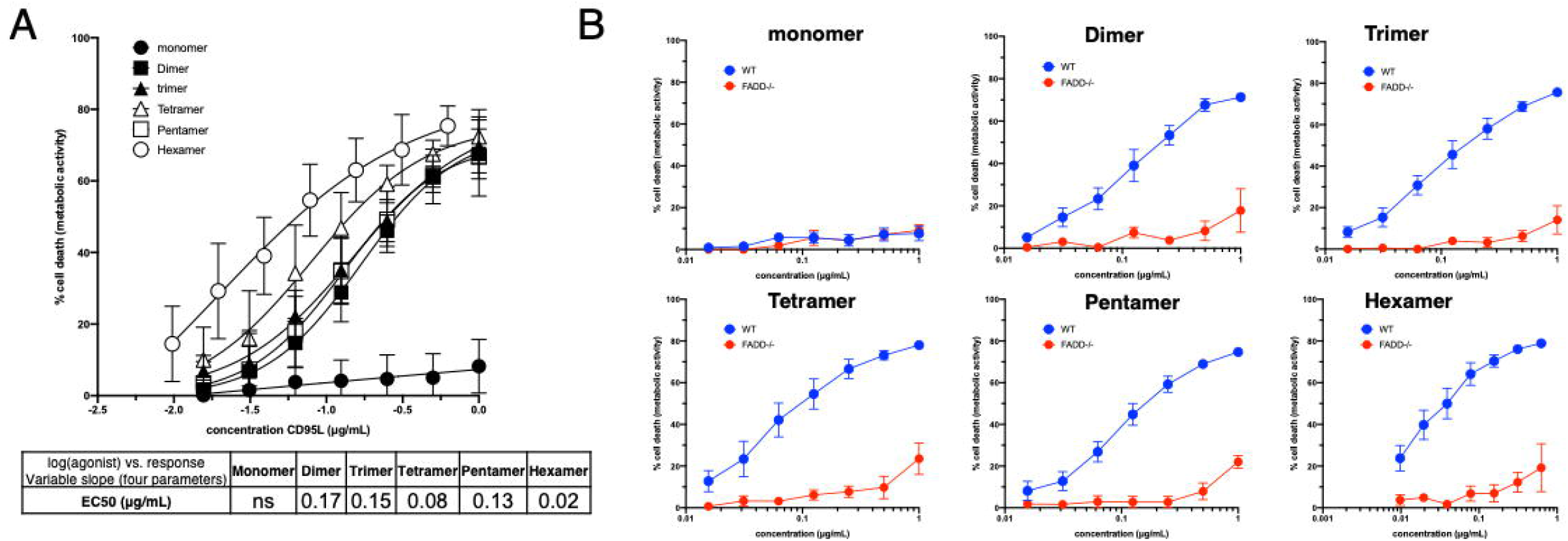
Concatemers of CD95L and induction of apoptosis. **A**. CD95L constructs secreted by HEK/293T cells were dosed by ELISA and their cytotoxic activity was evaluated by MTS assay using the CD95 sensitive cell line Jurkat. EC_50_ was assessed for the monomeric CD95L and the indicated concatemers. Data represent means and SD of three independently performed experiments. **B**. The cytotoxic activity of CD95L constructs was evaluated by MTS assay using the CD95 sensitive cell line Jurkat and its FADD-deficient counterpart, namely I2.1 cell line. Data represent means and SD of three independently performed experiments.

Surprisingly, although monomer and trimer were resolved as homo-trimers in BN-PAGE (Fig.1F), the former did not induce cell death in Jurkat cells, while the latter did (Fig.2A-B). These data suggested that either the minor self-association of trimer concatemer observed in BN-PAGE (Fig.1F) or these trimers exhibited two different trimeric conformations accounted for their difference in the induction of the apoptotic signal. We confirmed that the apoptotic signal induced by the concatemers relied on the adaptor protein FADD since FADD-deficient Jurkat cells did not undergo cell death when exposed to them (Fig.2B).

### CD95L concatemers and Ca^2+^ response

The CD95-mediated calcium response exerts a pleiotropic role, but we established that this signal is essential for cell migration of Th17 cells in autoimmune diseases such as lupus (14) and thereby, for the implementation of non-apoptotic functions (4,22). Interestingly, a dose effect performed with all concatemers revealed that cells exposed to trimeric and hexameric ligands underwent stronger CD95-mediated Ca^2+^ response as compared to those measured with the other ligands (Fig.3A). In agreement with our previous data (14,23), the metalloprotease-cleaved CD95L, which exhibited a homotrimeric structure (Fig.1F) also induced a strong Ca^2+^ response. Interestingly, although monomer, trimer and the metalloprotease-cleaved CD95L exhibited a homotrimeric structure (Fig.1F), the robust calcium signal observed with the metalloprotease-cleaved ligand and trimer concatemer (Fig.3B and 3C) was not phenocopied with the monomer (Fig.3A and 3C). CD95L can be cleaved by different metalloproteases and at different positions in its stalk region (24) such as ^113^EL^114^ (25), ^126^SL^127^ (3,4) or ^129^KQ^130^ (26,27) according to the literature. We previously observed that the metalloprotease-driven cleavage of CD95L in HEK/293T cells mainly occurs between amino acid residues ^126^SL^127^ (4) and thereby leads to the elimination of a part of the CD95L stalk region. Although the role of this stalk region remains to be elucidated, a long form of soluble CD95L containing the stalk region has been reported to induce cell death (19) and be responsible for the pathology severity in acute respiratory distress syndrome (20). Unlike the metalloprotease-cleaved CD95L, the monomer encompassed the stalk region and failed to trigger apoptosis (Fig.2A and 2B) and calcium response (Fig.3A and 3C), suggesting that beyond a putative role in the ligand aggregation, the stalk region could also affect the ligand conformation, and thereby alter CD95 activation. Nonetheless, it is noteworthy that the trimeric concatemer also possessed the stalk region and triggered both cell death (Fig.2A-B) and calcium response (Fig.3A & 3C) rendering difficult to apprehend the exact role of this juxtamembrane region.

**Figure 3.**
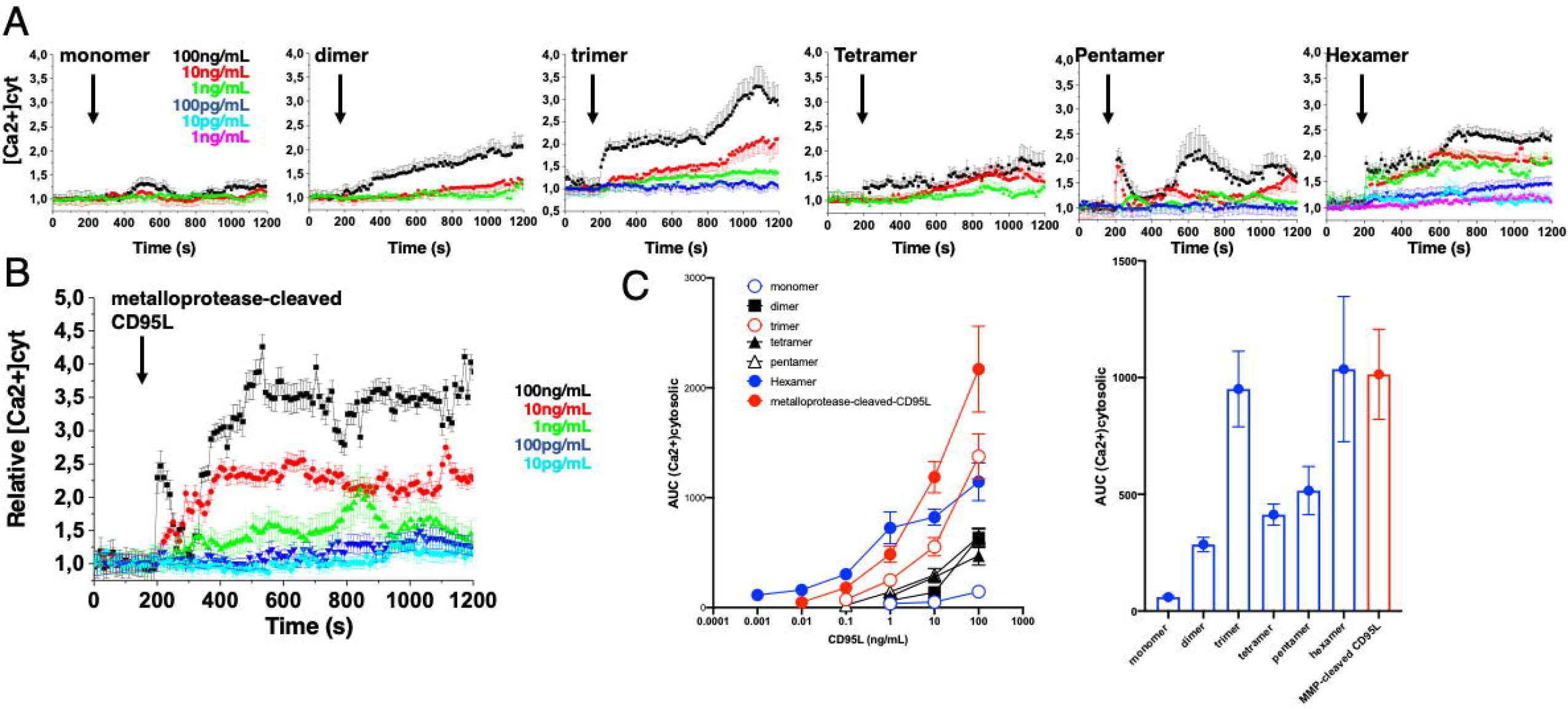
Concatemers of CD95L and induction of calcium response. **A**. The cytosolic calcium concentration ([Ca^2+^]_cyt_) in Jurkat cells exposed to the indicated concentration of CD95L was assessed using fura PE3-AM. Ratio values (R) were normalized to pre-stimulated values (R0) to yield R/R0 values (relative [Ca2+]). Data represent the mean □ ± □ SD. **B**. Same experiment as A with the metalloprotease-cleaved CD95L. **C**. *Left panel:* area under the curves (AUC) of the Ca^2+^ responses obtained in A and B are represented. *Right panel:* AUC of the Ca^2+^ response obtained in Jurkat cells exposed to the higher concentration of ligands (100 ng/mL).

In conclusion, this new set of ligands confirmed that the apoptotic signal relies on a aggregation threshold to be efficiently activated and hexamer is the best ligand to do it. On the other hand, the CD95-mediated calcium response relies on both conformation and stoichiometry because only trimer and hexamer (Fig.3A-C) can efficiently induce this response. Although the maintain of the stalk domain might explain why monomer, which self-associates as a trimer (Fig.1F), does not phenocopy the metalloprotease-cleaved ligand for the calcium response, this hypothesis has to be further investigated since the trimeric concatemer also encompasses this domain and efficiently implements apoptotic and non-apoptotic signals.

## Material and Methods

### Antibodies and reagents

Mouse monoclonal anti-FLAG M2 (#F3165) and anti-CD95 mAb CH-11 (#05-201) was from sigma-Aldrich (Sigma Aldrich Chimie, Saint-Quentin-Fallavier, France). Anti-CD95L antibody (G247-4, #556387) was acquired from BD Bioscience. MTS ([3-(4,5-dimethylthiazol-2-yl)-5-(3-carboxymethoxyphenyl)-2-(4-sulfophenyl)-2H-tetrazolium) (#G3581) was purchased from Promega (Charbonnières-les-Bains, France).

### Computer modeling

The chimeric FasL trimer was constructed from the CD95L:DcR3 complex (PDB: 4MSV,(18)). The receptor (DcR3) was eliminated and the N-terminal amino acid residues 103 to 143 were added in a completely extended form. A plausible conformation of the completed trimeric protein was obtained by molecular dynamics. Briefly, the built structure was solvated with 106500 water molecules and 16 chloride ions, in a 180 * 150 * 120 Å box for periodic boundary conditions. The whole protein was first kept frozen, while solvent and ions were optimized through 100 ps energy minimization. In a second step, only crystallized part of the protein was kept frozen for another 100 ps energy minimization. Then molecular dynamics was performed without any constraint for 1.4 μs. NPT conditions were used at 300 K and 1 bar, with a 2 fs timestep for near interactions, and 6 fs for distant interactions (RESPA integrator). All molecular mechanics experiments were performed with Desmond, as implemented in Maestro (Schrödinger LLC, San Diego, CA). Although each added N-terminal part was optimized individually and the 3 simultaneously, they converged to a partially folded structure, including 2 small α-helices involving residues 108-122. These helices were in close proximity to the loop bearing amino acids E185, K217 and S253.

### Cells

HEK/293T, parent Jurkat cell line (A3) and FADD-deficient (I2.1) counterpart was acquired from ATCC. HEK/293T cells were cultured in DMEM complemented with 8% FCS at 37°C in an atmosphere of 5% CO2. Jurkat cells were cultured in RPMI complemented with 8% FCS at 37°C in the presence of 5% CO2.

### Plasmids

Two pcDNA3.1(+) vector encoding CD95L-containing inserts were synthesized by Genecust (Boynes, France). These two vectors were designated vector 1 and 2. Vector 1 contained the CD95L monomer (amino acid 103 to 281 without stop codon) fused to (GGGGS)_2_ sequence according to the optimization of codon pair (28) with no stop codon. This CD95L-(GGGGS)_2_ sequence was surrounded by two restriction sites, BamHI in 5’ and BglII in 3’. Vector 2 contained the signal peptide (SP) of human albumin (29) synthesized in frame with the extracellular sequence of human CD95L (amino acid 102 to 281). Between albumin SP and CD95L sequences, we inserted three restriction sites (XhoI/BamHI/EcoRI) generating a LEGSE amino acid sequence. Vector 1 was digested by BamHI/BglII and the CD95L- (GGGGS)_2_ sequence was inserted in vector 2 digested by BamHI. Cohesive end of BamHI can be ligated with BglII site but the new ligation product is not cleavable by BamHI or BglII. Using this approach, we generated five concatemers of CD95L encoding two to six repetitions of human wild-type extracellular CD95L linked by the GGGS motif [(GGGS)_2_]. To generate the metalloprotease-cleaved CD95L, human full length CD95L sequence was synthesized by Genecust and inserted between EcoRI/BamHI in the MCS of pLVX-IRES-tdTomato vector. All vectors were verified by sequencing.

### Production of the soluble CD95Ls

Concatemers and full length CD95L-encoding vectors were transfected using calcium/phosphate method in HEK/293T cells. After 7 days of culture in serum-low Opti-MEM (Life technologies, Saint Aubin, France), secreted constructs or metalloprotease-cleaved CD95L were collected. Supernatants were ultracentrifuged for 1 h (100 000 g) to eliminate exosome contamination. Soluble CD95L concentration was quantified by ELISA.

### CD95L ELISA

Soluble CD95L was dosed by ELISA (#ab45892) following the manufacturer’s instructions (Abcam, MA, USA).

### Immunoblotting

Western-blots of HEK supernatants were performed as previously described (14). Briefly, protein concentration was determined by the bicinchoninic acid method (Pierce, Rockford, IL, USA) according to the manufacturer’s protocol. 30 μL (SDS-PAGE) and 13 μL (BN-PAGE) of supernatant were loaded per lane, resolved by BN- or SDS-PAGE gel and transferred to a PVDF membrane (Invitrolon™ PVDF Filter Paper Sandwich, #LC2005). Non-specific binding sites were blocked by incubating membranes for 60 min with TBST-Milk(M) (50 mM Tris, 160 mM NaCl, 0.05% (v/v) Tween 20 and 5% (w/v) dried skimmed milk at pH 7.4) and later incubated overnight at 4 °C with the corresponding primary antibody. For visualization of bound primary antibodies, membranes were washed with TBST, incubated with an appropriate peroxidase-labeled secondary antibody (anti-mouse IgG1-HRP #1070-05, Southern Biotech, Birmingham, USA) for 30 min. Membranes were revealed by enhanced chemiluminescence using Immobilon® Forte Western HRP Substrate (Millipore, #WBLUF0500) on Bio-Rad ChemiDoc™ Touch Imaging System.

### Blue Native-Polyacrylamide Gel Electrophoresis (BN-PAGE)

Protein constructs were subjected to native electrophoresis on NativePAGE™ Novex® Bis-Tris Gel system following manufacturer’s instructions (ThermoFisher). Briefly, supernatants (13 μl) were prepared using the NativePAGE™ sample Prep Kit (ThermoFisher, #BN2008) and loaded on 4-16% Bis-Tris NativePAGE™ gels (ThermoFisher, #BN1002BOX). Proteins were transferred to PVDF membranes (Invitrolon™ PVDF Filter Paper Sandwich, #LC2005) using NuPAGE® Transfer Buffer (ThermoFisher, #NP0006). Membranes were blocked with TBST-M for 1 hour at room temperature. The anti-CD95L mAb (G247-4) was incubated in TBST-M overnight at 4°C with gentle agitation. Then, membranes were washed with TBST buffer and secondary HRP-coupled antibodies were applied. Membranes were developed by enhanced chemiluminescence detection system using Immobilon® Forte Western HRP Substrate (Millipore, #WBLUF0500) on Bio-Rad ChemiDoc™ Touch Imaging System.

### Cell Death Assay

Cell viability was evaluated using MTS assay following the manufacturer recommendation. Briefly, cells (5.10^4^) were incubated for 24 h in flat-bottom 96-well plates with the indicated concentrations of the apoptotic inducers in a final volume of 100 μL. Then, 20 μL of MTS solution was added for 4 h at 37 °C. Absorbance was then measured at 490 nm wavelength and subtracted with a reference wavelength of 630 nm to eliminate nonspecific absorbance using the EnSpire^®^ 2300 Multilabel Plate Reader reader (Perkin Elmer, Waltham, Massachusetts, USA).

### Imaging and intracellular calcium concentration ([Ca^2+^]i) determination

Single-cell [Ca^2+^]I imaging was performed ratiometrically, using FuraPE3-AM calcium dye, as described previously (30). Briefly, cells were loaded with 1 μM FuraPE3-AM at room temperature in Hank’s Balanced Salt Solution (HBSS) for 30 min. FuraPE3-AM exhibits limited compartmentalization in intracellular stores and is leakage resistant (31). The cells were centrifuged to eliminate the fluorescent probe, rinsed with HBSS and incubated for 15 min to complete de-esterification of the dye. The loaded cells are deposited on a glass coverslip placed in a recording chamber positioned on the stage an inverted epifluorescence microscope (Olympus IX70) equipped with a ×40 Uapo/340-1.15W water-immersion objective (Olympus). To minimize UV light exposure, 4×4-binning function was used. FuraPE3-AM was excited at 340 and 380 nm alternately, and the ratios of the resulting images were captured at 12-bit resolution and constant 10-s intervals by a fast-scan camera (CoolSNAP fx Monochrome, Photometrics). Each experiment was independently repeated 3 times, and for each experimental condition, we displayed an average of more than 20 single-cell traces. Data were processed using OriginPro 7.5 software (Origin Lab). Fluorescence ratio (F_ratio_ 340/380, F) intensity changes were normalized to the initial fluorescence value F_0_ and expressed as F/F_0_ (relative [Ca^2+^]_cyt_). The calcium traces were quantified by determining associated areas under the curve (AUC) using Origin-Pro 7.5 software (OriginLab).

## Supporting information

Raw data wb

## Acknowledgments

This work was supported by INCa PLBIO (PLBIO 2018-132), ANR (ANR-22-CE15-0038-02) and Fondation de France (Price Jean Valade).

