## Supplementary figures and images for "CD95L concatemers highlight difference in the manner CD95-mediated apoptotic and non-apoptotic pathways are triggered"

### Raw data wb

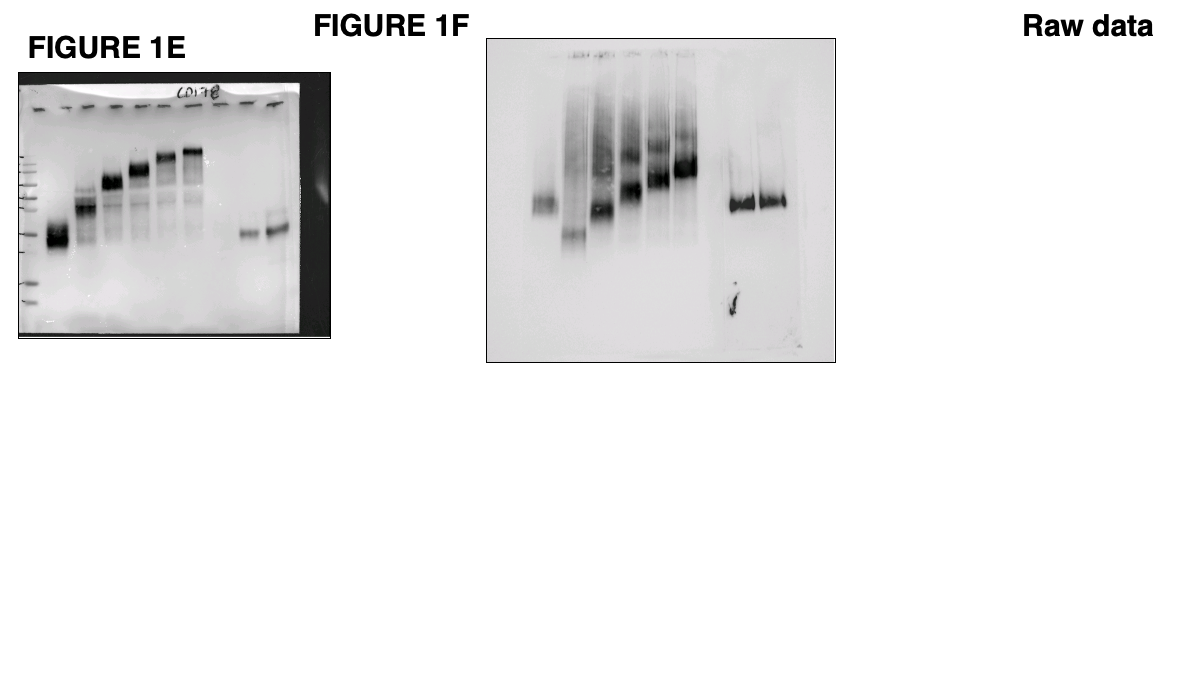
